# Systematic Engineering of Virus-Like Particles to Identify Self-Assembly Rules for Shifting Particle Size

**DOI:** 10.1101/2022.08.31.506130

**Authors:** Bon Ikwuagwu, Emily Hartman, Carolyn Mills, Danielle Tullman-Ercek

## Abstract

Virus-like particles (VLPs) are promising scaffolds for biomaterials as well as diagnostic and therapeutic applications. However, there are some key challenges to be solved, such as the ability to engineer alternate sizes for varied use cases. To this end, we created a library of MS2 VLP variants at two key residues in the coat protein which have been implicated as important to controlling VLP size and geometry. By adapting a method for systematic mutagenesis coupled with size-based selections and high-throughput sequencing as a readout, we developed a quantitative assessment of two residues in MS2 coat protein that govern the size shift in MS2 VLPs. We then applied the strategy to the equivalent residues in Qβ VLPs, an MS2 homolog, and demonstrate that the analogous pair of residues are also able to impact VLP size and shape. These results underscore the power of fitness landscapes in identifying critical features for assembly.

## Introduction

Protein assemblies are found everywhere in biology and industry and serve important natural functions. Virus-like particles (VLPs) are a self-assembling subset of such structures and garner particular interest due to their ease of production coupled with their encapsulating geometry. These features lend themselves to use in diverse applications such as enzyme spatial organization [1], therapeutic delivery [2–5], imaging [6], and biomaterial scaffolding [7]. Much effort has been directed towards understanding how self-assembly works [8, 9] for the further implementation of these useful protein assemblies. Despite the tremendous progress in predicting proper protein folding [10], the ability to alter quaternary structures to a specific design for an application remains lacking.

Viruses and their associated VLPs are an excellent model for protein self-assembly because they often 1) express well and in high titers, 2) are straightforward to purify, 3) harbor encoding nucleic acid sequences to link genotype to the resulting structures, and 4) form native structures with relatively high stability. Thus, single amino acid variants of their coat proteins can yield stable alternative structures. For example, a double variant of the tobacco mosaic virus (TMV) coat protein (CP) yielded a “nanodisk”[11] assembly instead of the standard rod assembly. Single variants in the CP of the bacteriophage Qβ yielded a size distribution of assemblies that included smaller, wild-type (WT)-sized, and longer prolate VLPs [12, 13]. A single amino acid variant in bacteriophage MS2 CP [S37P] (“Mini MS2”) [14] maintains nearly identical protein sequence, and has remarkably similar secondary and tertiary structure to the parent CP; however, the quaternary structure changes dramatically. Mini MS2 forms a 17 nm T=1 VLP instead of the 27 nm T=3 geometry WT MS2 VLP [14], such that the hexameric faces are eliminated in Mini VLP. Crystal structure analysis of both Mini and WT MS2 VLPs revealed different interdimer side chain interactions at residue 36 between WT and Mini MS2 VLPs, leading to a hypothesis that the change in interdimer interactions at residues 36 were a contributing factor for this shifted geometry. This shift illustrates how a single amino acid variation can access a large shift in the assembly mechanics of a protein complex, and further underscores the utility of the MS2 VLP as a model system.

Protein fitness landscapes offer the ability to comprehensively study the effect of a mutation in the amino acid sequence to a defined “fitness” metric used to generate the landscape. This defined fitness metric may be enzyme activity, binding, infectivity, or fluorescence, among other options [15]. We previously developed a method called SyMAPS (systematic mutagenesis and assembled particle selection) [16], in which a library of single amino acid mutations are generated in a VLP CP, followed by size-exclusion chromatography (SEC) to select for the ability to form icosahedral VLPs, the fitness applied in this context. Mutations are identified as present or absent following the selection by taking advantage of the native assembly dynamics of MS2. Specifically, the mRNA that encodes for a specific mutant is likely to be encapsulated and interacting with the interior of the VLP similarly to how the MS2 genome interacts with the native capsid [17]. Thus, after SEC and VLP extraction, mRNA is extracted from intact VLPs that encode for assembled particles. Reverse-transcription (RT-PCR) is used to transcribe this RNA into DNA, allowing high-throughput sequencing (HTS) on the assembly selections alongside the starting library plasmids. Fitness is quantified by calculating the ratio of HTS reads from the assembly selection to those of the starting plasmid library, enabling identification of variant MS2 VLPs that likely assemble. SyMAPS has since been adapted to study permitted residues and modifications at the N-terminus, a double mutation epistatic library at the pore[18], and a library containing all three-residue insertions into the MS2 FG Loop[19].

Here, we adapted SyMAPS to create a two-dimensional apparent fitness landscape (2D-AFL), which is an AFL for two mutations of the CP. We set out to assess if the side chain interaction differences we see in Mini MS2 can be used as a basis for uncovering assembly rules for VLPs with the smaller-sized phenotype. Our library consisted of 441 members, with degenerate NNK codons cloned into residues 36 and 37 in MS2 to encode for all 20 amino acids and a stop codon at both positions. (Figure 1).

**Fiugre 1:**
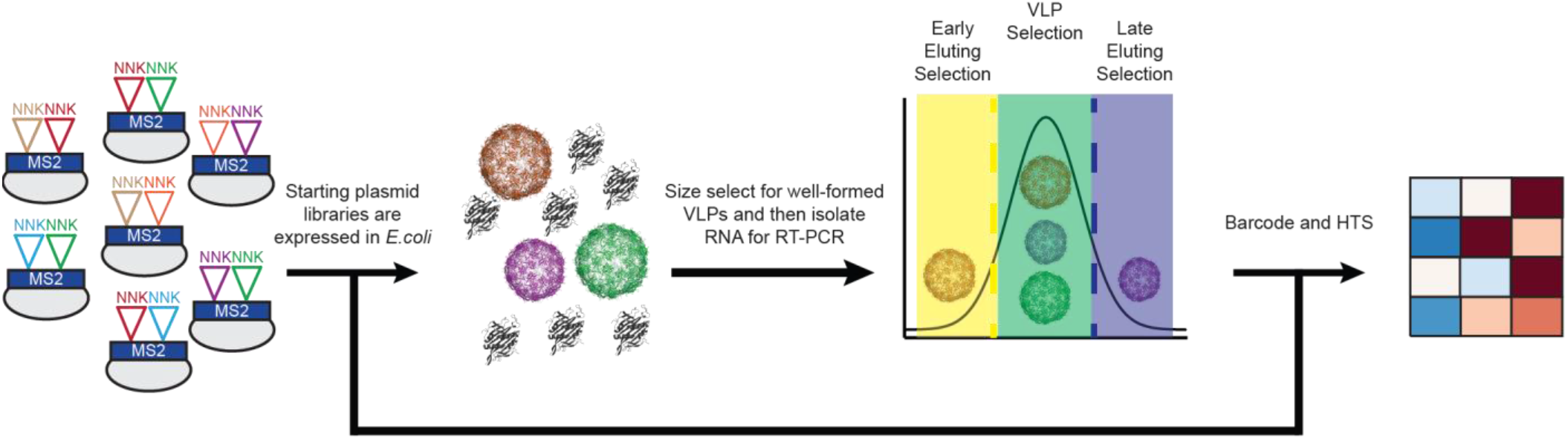
Modified SyMAPS protocol for MS2 36/37 NNK^2^ Library. The NNK codon = (Any)(Any)(G/T), so the resulting NNK2 Library has 441 members. Library is expressed, purified, and sequenced to generate an apparent fitness landscape.

Our approach takes advantage of the assembled particle selection step to separate smaller (e.g., Mini MS2-sized) VLPs from larger (e.g., MS2 WT-sized) particles. In this scheme, smaller diameter VLPs will elute in a distinctly later peak than the WT sized MS2 VLPs (Figure 1), offering the opportunity to identify the important contributions to the formation of each type of particle. We find that our hypothesis that residues 36 and 37 are important for Mini MS2 formation is supported. When mutations in this region are applied to an MS2 homolog Qβ, we see a similar smaller sized VLP phenotype emerge.

## Materials and Methods

### Strains and Media

Strains used in this study include *Escherichia coli* DH10B MegaX T1r DH10B *Escherichia coli* (ThermoFisher Scientific, Cat# C640003), which were used for all library experiments, and DH10B chemically competent cells produced in-house, which were used for expression of individual variants of interest. Overnight inoculums from a single colony were grown for 16-20 h at 37 °C shaking at 225 RPM in Luria Broth (LB)-Miller media (Fisher Scientific, Cat# BP1426-2) with chloramphenicol at 34 mg/L.

### Library Generation

To generate libraries with two amino acid mutations in MS2 and Qβ CPs, we modified a library preparation strategy used in prior studies to generate variant MS2 VLP libraries [14, 16, 17, 19]. The strategy, developed in the Bolon lab as EMPIRIC cloning [22], uses a plasmid with a self-encoded removable fragment (SERF) surrounded by inverted BsaI restriction sites. With this setup, BsaI digestion simultaneously removes both SERF and BsaI sites. These plasmids are termed “entry vectors”. The SERF in the entry vectors used in this study encodes constitutively expressed GFP to permit green/white colony screening. The MS2 36/37 NNK^2^ library was inserted into a previously generated entry vector of this design[16]. We created a new entry vector for the Qβ 40/41 NNK^2^ library via Gibson cloning, replacing residues 28 −54 with a constitutively active GFP flanked by BsaI sites.

The fragment that replaces the SERF in the entry vectors contains the double mutants for the 441-member insertion libraries. These fragments were synthesized using overlapping forward and reverse single-stranded DNA oligonucleotide primers, which were purchased through Sigma-Aldrich. Each primer was resuspended in water, pooled, and diluted to a final concentration of 50 ng/μL. The reverse strands were filled in by overlap extension (10 cycles of PCR). The resulting double-stranded DNA fragments were purified using a PCR Clean-up Kit (Promega, Cat# A9282). The purified DNA was then cloned into the entry vectors in place of the respective SERF using Golden Gate techniques [14, 16, 17, 19]. Briefly, a cycling Golden Gate method was performed that shifted between 37 °C for 2 minutes and 16 °C for 5 min for 24 cycles followed by 50 °C for 5min and 80 °C for 5 min. The resulting Golden Gate assemblies were desalted on membranes (Fisher Scientific# VSWP02500) for 5 min. Then MegaX DH10B *E. coli* was transformed with the resulting desalted Golden Gate assemblies and recovered for 1 h at 37 °C with shaking at 225 RPM. The recovered electroporated cells were plated on large (245 × 245 × 20 mm, #7200134, Fisher) LB-agar plates containing 34 mg/L chloramphenicol and allowed to grow overnight at 37 °C. To confirm that each unscreened, naïve library had at least three times the theoretical library size, we plated and counted 1:100 dilutions of recovered, transformed cells. This process was done in biological duplicate for the MS2 36/37 library and biological triplicate for the Qβ 40/41 library at every step.

### VLP Library Expression and Purification

Colonies for each library replicate were individually scraped from their plates into 5 mL of 2×YT with 5 mL 2×YT with 34 mg/L chloramphenicol and grown for 2 h at 37 °C with shaking at 225 RPM. Glycerol stocks were made with equal OD equivalents saved in each tube. Each replicate was then subcultured into 1 L of 2×YT with 34 μg/mL chloramphenicol. The cultures were grown to an OD_600_ of 0.4-0.6, and then expression was induced with 0.02% (w/v) L-(+)-arabinose. The NNK^2^ double mutant libraries were expressed at 37 °C with shaking at 225 RPM overnight (16-20 h). The cells were then harvested by centrifugation at 4,800 × g for 10 min, resuspended in 10 mM sodium phosphate, 200 mM sodium chloride, pH 7.2 buffer, and sonicated at 50% amplitude, pulsing 2 s on and 4 s off for a total of 10 min of “on” time (Fisher Scientific, catalog no. FB120A220, probe CL-18). Note that from harvesting onward the libraries were kept at 4 °C. The libraries were precipitated overnight with 50% (w/v) ammonium sulfate and collected by centrifugation at 17,000 × g for 10 min. Pelleted precipitates were then resuspended in 10 mM sodium phosphate, 200 mM sodium chloride, pH 7.2 buffer and 50% (w/v) ammonium sulfate precipitated for at least one hour. Samples were then collected by centrifugation at 17,000 × g for 10 min resuspended in 10 mM sodium phosphate, 200 mM sodium chloride, pH 7.2 buffer and syringe filtered (Polyethersulfone (PES) 0.45 μm, Fisher Scientific catalog no. 05-713-387).

### FPLC SEC (Assembly Selection)

VLPs were purified on an Akta Pure FPLC system via SEC with a HiPrep Sephacryl 16/60 S-500 HR (GE Healthcare Life Sciences, Cat# 28935607) column. Isocratic flow was used to elute, with a 10 mM sodium phosphate, 200 mM sodium chloride, pH 7.2 buffer. Fractions expected to contain MS2 CP were collected for further analysis (fractions from 0.47 – 0.7 column volumes (CV), representing the fractions that harbor WT and/or Mini MS2 VLPs). To identify double mutants with different sizes, fractions from 0.47 – 0.5 CV were pooled as the “early fraction” and separately analyzed, as were fractions from 0.67 – 0.7 CV as the “late fraction”. Similarly, fractions containing Qβ CP were collected for further analysis based on where WT Qβ is observed to elute under the same conditions (fractions from 0.47 – 0.7 CV). To identify double mutants with different sizes, the same CV fractions were collected for the Qβ “early fraction” and “late fraction.”

### Sample Preparation for High-Throughput Sequencing

Plasmid DNA was extracted from frozen glycerol library aliquots using a Zyppy Plasmid Miniprep Kit according to manufacturer instructions (Zymo, Cat# D4036). After performing assembly selections on the double mutant libraries, RNA was extracted from the samples taken from SEC using previously published protocols [14, 16, 17, 19]. Briefy, TRIzol (Thermo Fisher Cat# 15596026) was used to homogenize samples, followed by chloroform addition. The sample was separated by centrifugation into aqueous, interphase, and organic layers. The aqueous layer, which contained RNA, was isolated, and the RNA was then precipitated with isopropanol and washed with 70% ethanol. RNA was then briefly dried and resuspended in RNase-free water. cDNA was then synthesized using Superscript III first-strand cDNA synthesis kit from Life Technologies (catalog no. 18080051, random Hex primer). cDNA and plasmids were both amplified with two rounds of PCR to add barcodes (10 cycles for MS2, 13 cycles for Qβ) and the Illumina sequencing handles (8 cycles), respectively, following Illumina 16S Metagenomic Sequencing Library Preparation recommendations. Libraries were combined and analyzed by 300 PE MiSeq in collaboration with the Lucks Lab at Northwestern. Reads in excess of 2 million passed filter, and an overall Q30 > 75%.

### High-throughput Sequencing Data Analysis

Data were trimmed and processed as previously described[16, 19, 20] with minor variation. Briefly, data were trimmed with Trimmomatic [21] with a 4-unit sliding quality window of 20 and a minimum length of 30.

### 2D-AFL calculations

Trimmed high-throughput sequencing reads were analyzed using Python programs written in-house and uploaded onto Github. Briefly, reads encoding for the first 57 residues of the MS2 or Qβ CP were isolated, and the number of mutations per read was calculated. Reads with zero mutations (WT reads) or greater than two mutations were both removed. In reads with two mutations, the two non-wild-type codons were identified and counted. In reads with one mutation, the mutated codon was tallied in combination with every wild-type codon. Non-wild-type codons that did not end in G or T were also eliminated as these were not encoded by NNK. Codons were then translated into amino acids. These calculations were repeated for all experiments to generate abundances before and after each assembly selection. Relative percent abundances were calculated as previously described [14]. Briefly, the grand sum, or the sum of all counts at every combination at the two residues of interest, were calculated. We next divided the matrix by its grand sum, generating a matrix of percent abundances. These calculations were repeated for each biological replicate of plasmid, VLP fraction, early fraction, or late fraction, generating eight Percent Abundance matrices for the MS2 36/37 library and twelve Percent Abundance matrices for Qβ 40/41 library. We calculated relative percent abundances by dividing the Percent Abundance for the selected library compared by the percent abundance for the plasmid library for each replicate.

We calculated the mean across two replicates for MS2 and three replicates for Qβ. All Nan (null) values, which indicate variants that were not identified in the plasmid library, were ignored. Scores of zero, which indicate variants that were sequenced in the unselected library but absent in the VLP library, were replaced with an arbitrary score of 0.0001. We calculated the log10 of the Relative Percent Abundance array to calculate the final array for each replicate. Finally, we calculated the average apparent fitness score (AFS) value for each amino acid combination by finding the mean value for every combination across replicates.

### Mini Forming Propensity Apparent Fitness Score Calculation

2D-AFLs were generated using the early fraction and late fraction from the FPLC SEC assembly selection barcoded DNA and the starting plasmid library barcodes. The MFP AFS is the ratio of the 2D-AFL from the late fraction over the 2D-AFL from the early fraction.

### HPLC SEC

VLP variants were analyzed on an Agilent 1290 Infinity HPLC with a YARA-4000 SEC column (5 μm, 2000 Å, 7.8×300 mm) with isocratic flow of 10 mM sodium phosphate, 200 mM sodium chloride, pH 7.2 buffer. Fractions were collected at the characteristic elution peak for WT MS2 and Qβ VLPs (7.5 −8.5 min) and Mini-sized VLPs (9.4-10.6 min).

### Relative Size Bias

Area under the WT and Mini HPLC characteristic peaks are calculated for each biological replicate of an individual variant using the trapezoid rule. The relative size bias is the ratio of the area under the characteristic Mini peak over the sum of the areas of the WT and Mini characteristic peak. A value of one means there was no area under the WT characteristic peak while a value of 0.5 means that the area under the WT and Mini peaks are equivalent.

### Individual Variant Cloning

Individual variants were cloned using a variation on the method described earlier. Briefly, overlap extension PCR yields a double stranded fragment that spans the length of the missing 26-codon region in the entry vector. Each fragment was cloned into the entry vector using standard Golden Gate cloning techniques. Cloned plasmids were transformed into chemically competent DH10B cells. Individual clones were sequence confirmed via Sanger Sequencing (Quintara Biosciences) prior to expression. All plasmids were stored in DH10B cells and plasmid names described in Supplemental Table 1 refer to strain transformed with the plasmid.

### Individual Variant Expression

Selected variants were individually expressed in 50 mL cultures of 2×YT following the same procedure as for the libraries. The cultures were pelleted, resuspended in 10 mM sodium phosphate, 200 mM sodium chloride, pH 7.2 buffer, lysed by sonication, precipitated with 50% w/v ammonium sulfate, and evaluated by HPLC for VLP assembly.

### Transmission Electron Microscopy (TEM)

VLP samples purified by HPLC were collected and spin concentrated with a centrifugal filter with a 100 kDa molecular weight cutoff (Millipore Sigma, catalog no. UFC510024) at 5,000 G for 10 min. All VLP samples greater than 10 mg/mL were diluted down to 10 mg/mL for TEM visualization. VLP samples were fixed with 2% (v/v) glutaraldehyde in water solution on 400 mesh Formvar-coated copper grids (EMS Cat#FF400-Cu). After fixation, grids were washed with MilliQ™ pure water and stained with 1% (w/w) uranyl acetate in water. Grids were dried and stored prior to imaging. Images were acquired on a JEOL 1230 transmission electron microscope with a Gatan 831 bottom-mounted CCD camera with a 200 kV accelerating voltage. TEM Images were contrast-adjusted and cropped using ImageJ [22]. For VLP sizing, images were scale-corrected based on the instrument used to collect the images. The oval tool was used to manually trace an ellipse surrounding VLPs. The diameter of the ellipse, corresponding to the diameter for the VLP, was recorded. Further data analysis was carried out using Microsoft Excel or Python.

## Results and Discussion

### Generating a 2D-AFL for MS2 36/37

Based on a comparison of previously published crystal structures of MS2 and Mini MS2 [14], we hypothesized that residues 36 and 37 in MS2 are crucial to forming Mini MS2 VLPs due to the change in sidechain interactions, such as hydrogen bonding, at residue 36. The change from the MS2 WT VLP is a result of the proline mutation at residue 37 shifting the structure of the Mini MS2 CP. To test this hypothesis, we created a 441-member library in which residues 36 and 37 were encoded as NNK to ensure we tested all possible amino acids simultaneously at these two positions. The mutations were introduced into a pBAD-inducible vector. *E. coli* cells harboring these vectors were collected into a single culture, grown, and induced for expression of all library members. To identify combinations of residues that conferred Mini MS2-sized particles as well as WT sized particles, we selected for assembled particles using fast protein liquid chromatography (FPLC) SEC. To do so, we recovered fractions from the characteristic VLP elution peak and from both the leading (earlier) portion of the peak and the lagging (later) portion of the VLP peak. Given that larger particles elute earlier than smaller particles with SEC, we reasoned that these selections would be biased toward larger and smaller particles, respectively (Figure 1). The overall VLP selection included the elution range starting at the early fraction and ending at the larger fraction. To identify the sequences encoding each type of particle, we first extracted RNA from all three of the collected assembly fractions, because mRNA encoding an assembly competent variant MS2 VLP is encapsulated within the VLP during the assembly of that particle within the host cell[23–25]. We then performed RT-PCR to generate the DNA that encoded the mutations from our assembled VLP variant. We barcoded these samples and our starting plasmid library for each biological replicate. After high-throughput sequencing, the 2D-AFL was then generated from an AFS [14] for each member of the library, which is calculated by taking the log10 of the ratio between the relative abundance of the plasmid library over the relative abundance of the VLP selection fraction reads. An artificial AFS of −4.0 is assigned to mutants that are present in the plasmid library but not sequenced in the VLP assembly fractions. Based on our prior work, we find that an AFS > 0.2 indicates assembly competent VLPs and an AFS < −0.2 indicates assembly deficient VLPs [14, 16, 17, 19].

### Assessing the Quality of the MS2 36/37 2D-AFL

To explore the coverage of the 2D-AFL, defined as how many members of the library are present, we analyzed the counts of each possible member in the biological replicates of the plasmid library. The starting plasmid library is the basis for any fitness calculation, so if a double mutant is missing in the plasmid library, then no fitness data can be computed for that double mutant. Approximately 99.9% of the library was present in the biological replicates of the plasmid library reads, with missing mutants colored yellow and striped in our AFL (Figure 2A). Of the eight missing double mutants, three were designed to encode the TAG stop codon, so we do not expect those variants to form assembly competent VLPs. With a significant portion of the plasmid libraries present for expression, we are confident that the analysis we derive from the 2D-AFL reflects the nature of the sequence space of possible mutations at residues 36 and 37 of the MS2 CP.

**Figure 2:**
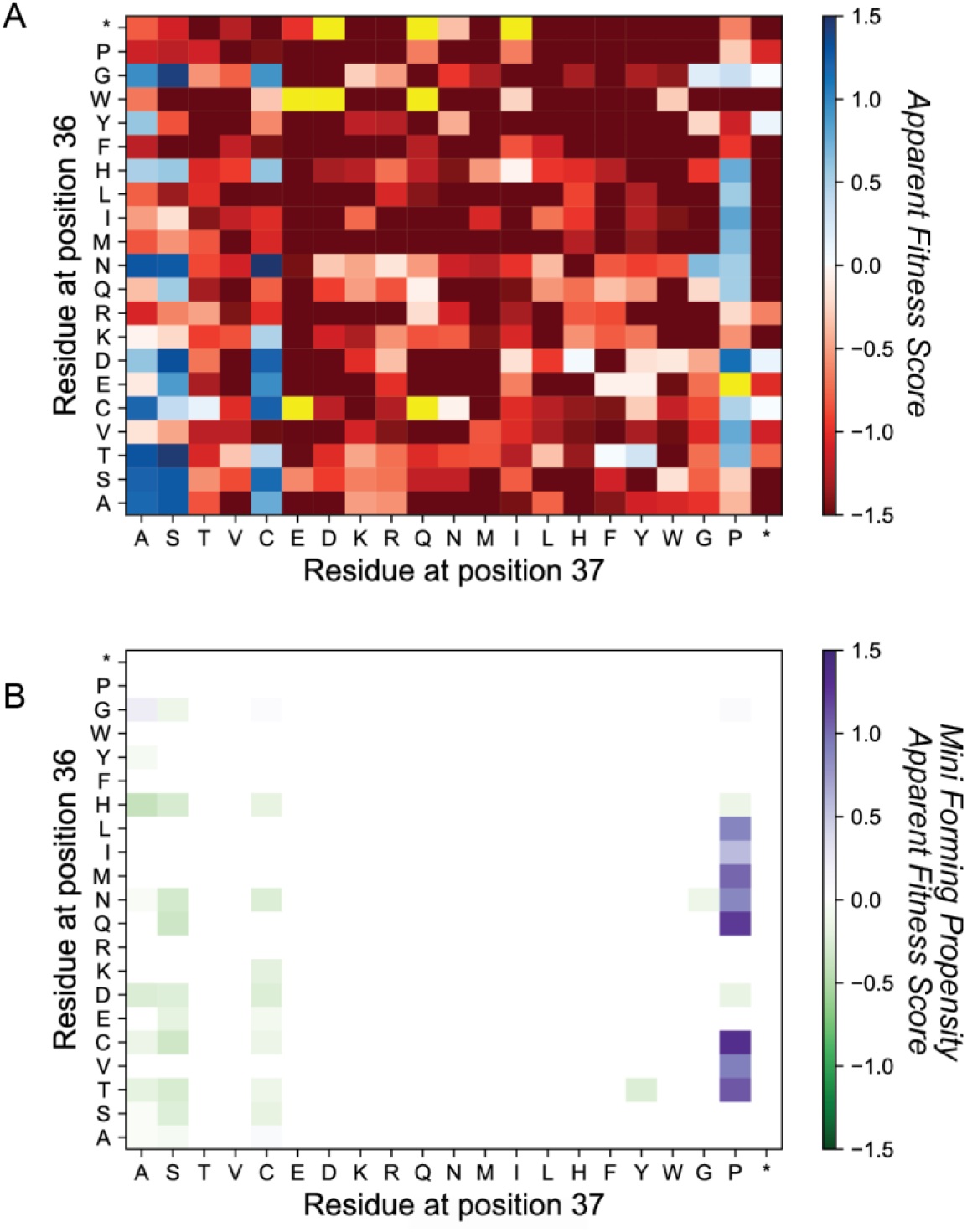
2D-Apparent Fitness Landscape of MS2 36/37 Library. A) 2D-AFL of the VLP selection library. Blue indicates double variants were enriched after assembly selection and red indicates double variants that were less abundant after selection. Yellow indicates missing values. B) Composite 2D-AFL where Mini Forming Propensity Apparent Fitness Scores are overlayed double variant combinations whose AFS > 0.2. Purple indicates variants were more abundant in the later eluting fractions vs the earlier eluting fractions. Green indicates variants were more abundant in the later eluting fractions vs the earlier eluting fractions.

Analysis of the 2D-AFL was next performed to determine whether scores at nonsense mutations were consistent with their expected value. Nonsense mutations are mutations in which a stop codon is encoded and thus should not form full length protein, so we expected an AFS < −0.2 for any variant encoding at least one nonsense mutation. 38/41 nonsense mutations were present in our plasmid library and the average AFS for these mutations are −2.2, consistent with what we expected for non-assembling mutants. Only four of the possible nonsense mutations have an AFS > 0, but those variants are all less than 0.2, and are thus below the cutoff for VLP assembly. Approximately 66% of nonsense mutants are screened out of the VLP library completely, which is indicated by an AFS of −4.

### Identifying Trends and Validating the MS2 36/37 2D-AFL

Using our heuristic AFS > 0.2 as a cutoff for assembly, we observed that assembly competent VLPs have either an alanine, cysteine, proline, or serine at position 37 (Figure 2A). To validate that an AFS > 0.2 is predictive of icosahedral VLPs with this library, we constructed a subset of double mutants with an alanine, cysteine, or proline at position 37 and used HPLC SEC to measure assembly. Assembly is evidenced by a higher A_260_ peak than A_280_ peak at a given position, so we considered a particle to be assembled if A_260_ > A_280_ either at the characteristic WT VLP elution time, 7.5 −8.5 min, or at the characteristic Mini VLP elution time, 9.4-10.6 min [14]. For MS2 WT (CP[NS], AFS = 1.3) and Mini MS2 (CP[N36/S37P], AFS = 0.54), the HPLC SEC assembly assay yielded expected peaks in the WT or Mini elution times respectively.

We next examined select members of the 2D-AFL with a focus on those that contain a flexible amino acid at residue 36 and assemble per our heuristic of an AFS > 0.2. First, we checked that three predicted assembling double mutants, MS2 CP[N36G/S37A], MS2 CP[N36G/S37C], and MS2 CP[N36G/S37P], formed particles with the flexible amino acid glycine at residue 36. We individually cloned each double mutant, leveraging our library’s Golden Gate “entry vectors” to clone the variants into an expression plasmid. We performed small scale 50 mL expressions of these three variants, purified with the same techniques as used with the library, and then assessed the VLP assembly via HPLC SEC. All three of these double variants elute at the expected MS2 WT elution time and have A_260_ > A_280_, even though some traces are broader than the MS2 WT traces (Supplemental Figure 1).

We next examined two assembly deficient members of the library, introducing a bulky tryptophan or another proline into residue 36. Our HPLC SEC-based assembly validation of MS2 CP[N36W/S37P] (AFS = −4) and MS2 CP[N36P/S37P] (AFS = −2.1) revealed that no peaks were present in any characteristic VLP elution ranges, consistent with the non-assembling phenotype we expected. (Supplemental Figure 2 A-B). We finally wanted to examine the assembly nature of library members for which we were unable to predict assembly from the AFS, such as MS2 CP[N36E/S37F] and MS2 CP[N36D/S37H], which both have an AFS = ~0. As assessed by our HPLC SEC-based method, these mutants did not form VLPs (Supplemental Figure 2 C-D), supporting our hypothesis that an AFS > 0.2 continues to represent VLP assembly for this 2D-AFL.

### Identifying Different Sized MS2 VLPs using the Mini Forming Propensity AFS

With our assays established and validated, we returned to our goal of understanding how mutations in residues 36 and 37 affect Mini MS2 formation propensity. To identify different sized MS2 VLPs using the SyMAPS approach, we simultaneously generated 2D-AFLs based on the later eluting fractions and earlier eluting fractions of the FPLC SEC assembly selection (Figure 1). We hypothesized that the ratio of the AFSs derived from the later eluting AFL over the AFSs derived from the early eluting AFL (Supplemental Fig 3 A-C) will provide us with a useful metric to identify potential smaller VLPs. We define this comparison metric as the “Mini MS2 Forming Propensity” (MFP) AFS. We further constrained the set to those variants that have an AFS > 0.2 in the assembly 2D-AFL. Our composite MFP 2D-AFL therefore includes only the double mutant-containing variants in the library that are likely to form Mini MS2 sized VLPs as shown in purple (Figure 2B).

The composite MFP 2D-AFL revealed 12 MS2 CP double mutants that potentially form Mini MS2 sized VLPs. Although MS2 CP[N36G/S37A], MS2 CP[N36G/S37C], and MS2 CP[N36G/S37P] MFP AFSs > 0, we had already determined these double mutants do not have peaks in the characteristic Mini VLP elution times (Supplemental Figure 1). Interestingly, 9/12 of the double mutants include the Mini MS2 CP [S37P] mutation. Moreover, 8/9 of the MS2 CP[S37P]-based double mutants have an AFS > 0.2 and an MFP AFS ≥0.2 (Figure 2B). The trend reaffirms the importance of a benchmark for a physical assembling phenotype being set at an MFP AFS ≥0.2, rather than MFP AFS ≥ 0, just as we observed with prior MS2 fitness landscapes.

HPLC SEC assembly validation revealed that members of the composite 2D-AFL with MFP AFS ≥0.2 form Mini MS2-sized VLPs (Figure 3A). We see a smaller peak with a higher A_260_ than A_280_, consistent with VLP formation, appearing in the later time frame of the characteristic WT elution peak for select double variants such as MS2 CP[N36I/S37P] and MS2 CP[N36T/S37P]. The presence of this smaller peak raised the hypothesis that other VLP assemblies were forming with these double variants.

**Figure 3:**
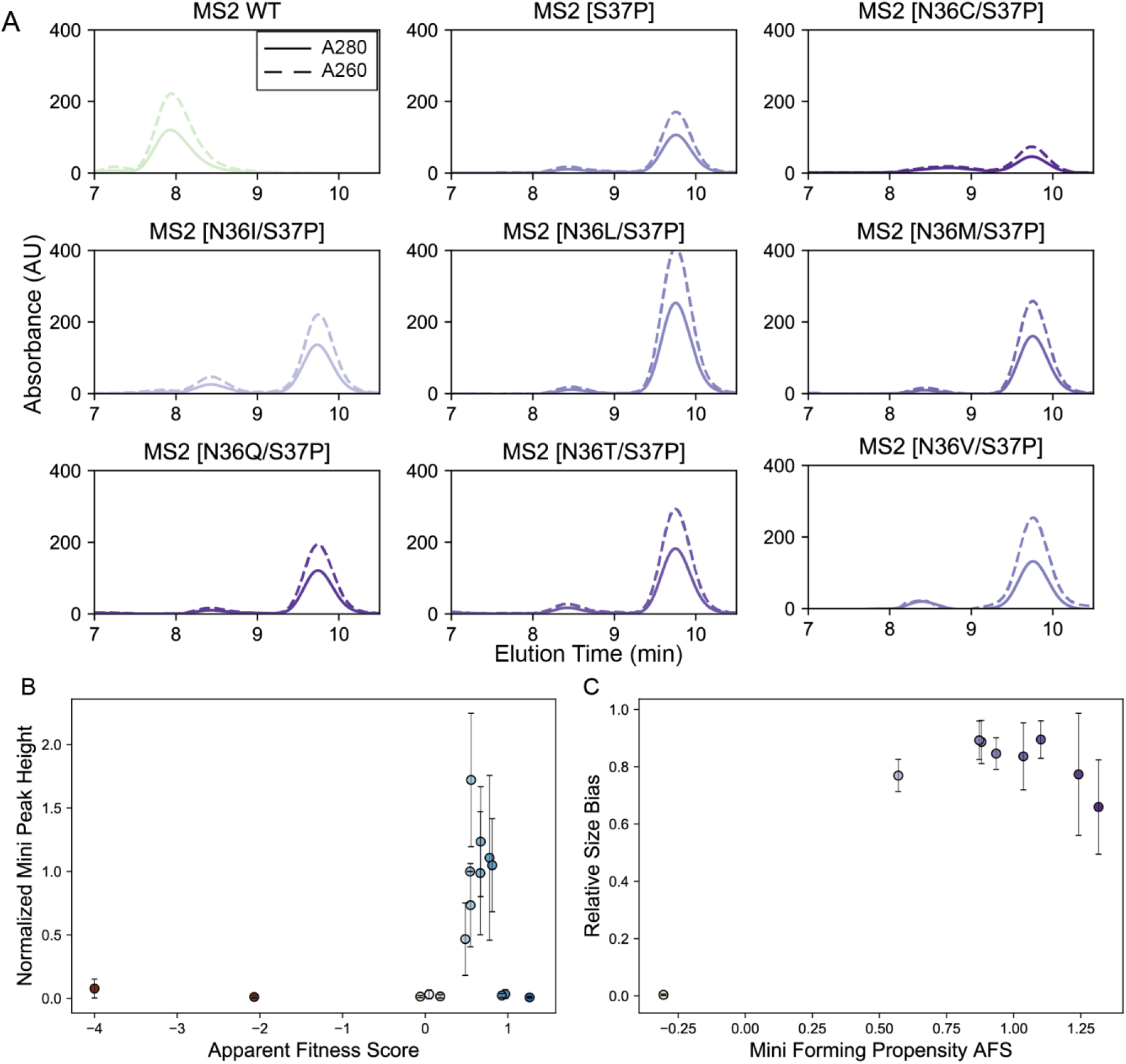
HPLC SEC Validation of MS2 36/37 2D-AFL. A) HPLC Traces of double mutants that have AFS >0.2 and Mini Forming Propensity (MFP) AFS >0.2. Traces colored by MFP AFS value. B) Normalized Mini Peak Heights plotted against the double variants AFS. Peak heights were normalized against MS2 CP [S37P] (n=3) C) Relative Size Bias plotted against MFP AFS

### Characterization of MS2 CP[S37P] Double Mutant VLPs

We next assessed if the different amino acid substitutions to residue 36 among the eight MS2 CP[S37P]-containing double variants influenced the extent of Mini MS2 VLP formation. For each individual double mutant, we analyzed the A_280_ traces, a proxy for protein concentration, from the characteristic Mini MS2 VLP peak. We calculated the maximum peak height for each double variant and normalized to the Mini MS2 value. Four double mutants, MS2 CP[N36L/S37P], MS2 CP[N36V/S37P], MS2 CP[N36I/S37P], and MS2 CP[N36T/S37P], had normalized Mini peak height values > 1, possibly forming more Mini MS2 VLPs than the original MS2 CP[S37P] (Figure 3B). For three of these (MS2 CP[N36L/S37P], MS2 CP[N36V/S37P], MS2 CP[N36I/S37P]), the non-native amino acid sidechains for the mutated residues do not have hydrogen bonding interactions such as the asparagine at residue 36[14] in Mini MS2, so the presence of Mini-sized MS2 VLPs was unexpected.

We next sought to understand if MS2 CP[S37P]-including variants formed homogeneous or heterogeneous VLP size populations. To do so, we analyzed the elution peak areas from the HPLC SEC assembly fractions compared to their MFP AFSs. The relative size bias is a metric that compares the maximum normalized peak area in Mini and WT elution time ranges. A relative size bias of one is analogous to Mini MS2 behavior, while a relative size bias = 0.5 would indicate an even split between Mini MS2 and MS2 WT peak areas (Figure 3C). MS2 WT has a relative size bias of 0.0038 and Mini MS2 has a relative size bias of 0.89 by this metric, giving us confidence that the metric would reflect different-sized VLP phenomena. Of the eight double variants predicted to form a Mini MS2-sized particle, MS2 CP[N36C/S37P], MS2 CP[N36I/S37P], and MS2 CP[N36T/S37P] have the lowest relative size bias scores at 0.65, 0.76, and 0.77 respectively. Of these three double mutants, MS2 CP[N36C/S37P] has an MFP AFS of 0.93, MS2 CP[N36T/S37P] has an MFP AFS of 1.1, and MS2 CP[N36I/S37P] MFP AFS is the lowest of the mutants predicted to form Mini MS2 VLPs with an MFP AFS of 0.567, which is still well above the 0.2 threshold. We identified MS2 CP[N36I/S37P] as a likely candidate for forming a heterogeneous population of both Mini and WT sized VLPs due to the combination of having a higher A_260_ than A_280_ in both the Mini MS2 and MS2 WT characteristic elution ranges, having the lowest relative size bias, and the lowest MFP AFS of our eight selected double variants.

To assess the possibility of other sizes or geometries arising in our samples, we decided to use TEM to study individual particles at higher resolution. To do so, we modified the HPLC SEC VLP assembly assay to collect fractions ranging from the characteristic WT to the characteristic Mini elution peaks. We combined the fractions into one sample, concentrated the sample, and then loaded the supernatant onto grids for TEM analysis. We performed this assay for Mini MS2 as a positive control for mini-size VLP assembly. We also examined MS2 CP[N36V/S37P] as a sample likely to form exclusively Mini-sized MS2 VLPs, and MS2 CP[N36I/S37P], which our various metrics indicated may form other sizes or geometries in addition to Mini-sized VLPs. All the variants chosen had visual evidence of Mini VLPs by TEM (Figure 4A-C). We see that in MS2 CP[N36I/S37P] we have a majority of Mini MS2 VLPs, but some evidence of WT sized VLPs (Figure 4, B and D).

**Figure 4:**
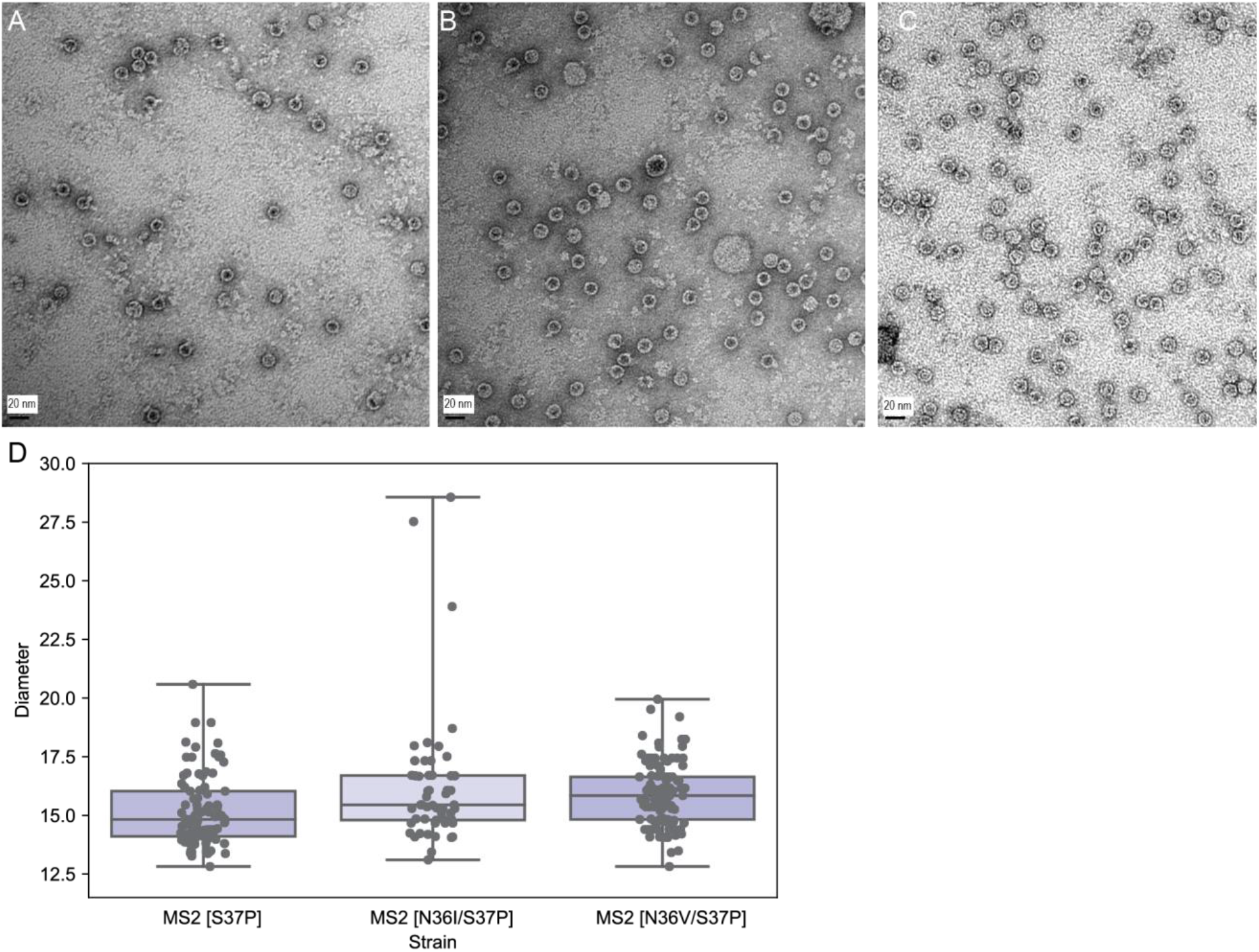
TEM Analysis of individual variants for Mini-sized MS2 VLPs. Black bar represents 20 nm. A) MS2 CP[S37P] B) MS2 CP[N36I/S37P] C) MS2 CP[N36V/S37P] D) Diameters of at least 50 VLPs quantified from TEM images are shown.

TEM image analysis was performed with at least 90 particles counted per double variant. TEM analysis on MS2 CP[N36I/S37P] characteristic WT elution fraction alone also showed that WT VLPs are forming (Supplemental Figure 4), so we were confident that MS2 CP[N36I/S37P] forms a heterogeneous population of MS2 VLPs.

The change in sidechain interactions due to the proline mutation at residue 37 likely contributes to MS2 VLPs forming smaller 17 nm particles. Analysis of the crystal structure of MS2 WT VLPs revealed that there are two hydrogen bonding interactions between native asparagine in residue 36 in a C/C dimer and native asparagine in residue 98 in the B conformation of an A/B dimer[14, 26]. These two hydrogen bonds were between the sidechain oxygen at residue 36 in the C conformation and backbone nitrogen at residue 98 in the B conformation and conversely the sidechain oxygen at residue 98 in the B conformation and the backbone nitrogen at residue 36 in the C conformation. In the Mini MS2 VLP crystal structure, the sidechain interactions between residues 36 and 98 changed to the sidechain nitrogen in the asparagine at residue 36 of one monomer and the backbone oxygen in the asparagine at residue 98 of an opposing Mini MS2 CP monomer. We hypothesized that this interaction difference, the single inter sidechain hydrogen bonding interaction, could lead to more flexibility in VLP assembly for Mini MS2 VLPs. We see that Mini MS2 CP have more flexibility in forming CP dimers, with Mini MS2 having five distinct conformations compared to the three conformations in WT MS2 CP. With the other known nitrogen containing amino acid, glutamine, at position 36, MS2 CP[N36Q/S37P], homogeneous size populations of Mini MS2 VLPs are formed. Threonine at residue 36 potentially hydrogen bonds with the side chain nitrogen at residue 98 and not the oxygen in this residue, yet we see MS2 CP[N36T/S37P] VLPs form a homogeneous population of Mini MS2 VLPs. Moreover, five of the eight double mutants we validated as forming Mini MS2 VLPs did not have any potential hydrogen bonding interaction at residue 36. To further explore the assembly dynamics of these double variant MS2 VLPs, Cryo-EM of specific variants should be performed, especially MS2 CP[N36Q/S37P], which has the highest probability of having similar side chain interactions of Mini MS2 VLPs and MS2 CP[N36I/S37P] yet appears to form a heterogeneous population with respect to size.

### Applying Design Heuristics for Mini MS2 VLPs to MS2 Homologs

We hypothesized that the heuristics we identified for MS2 are generalizable to other VLPs. There are many other bacteriophages that can form VLPs by overexpressing a coat protein, and a number of these are structurally similar to MS2 [26, 27]. We decided to study the CP of Qβ bacteriophage as a second model VLP, because it has a solved structure, forms VLPs, and is at least 20% similar in coat protein sequence to MS2. (Figure 5A). We rationally imposed a proline mutation into residue 40 of Qβ, the analogous position to residue 37 in MS2, to see if we could recreate the smaller sized VLP phenotype (Figure 5A). Both residue 40 in Qβ and residue 37 are in unstructured loop regions of their coat protein (Figure 5B-C). Therefore, even if there were less interaction with other amino acids at residue 40 in Qβ, we hypothesized that a proline mutation could shift the CP in similar ways to Mini MS2 CP. TEM visualization of Qβ CP[A40P] mutants demonstrate a heterogenous size population results from this mutation, which confers both WT Qβ-sized and smaller-sized VLPs (Figure 5E).

**Figure 5:**
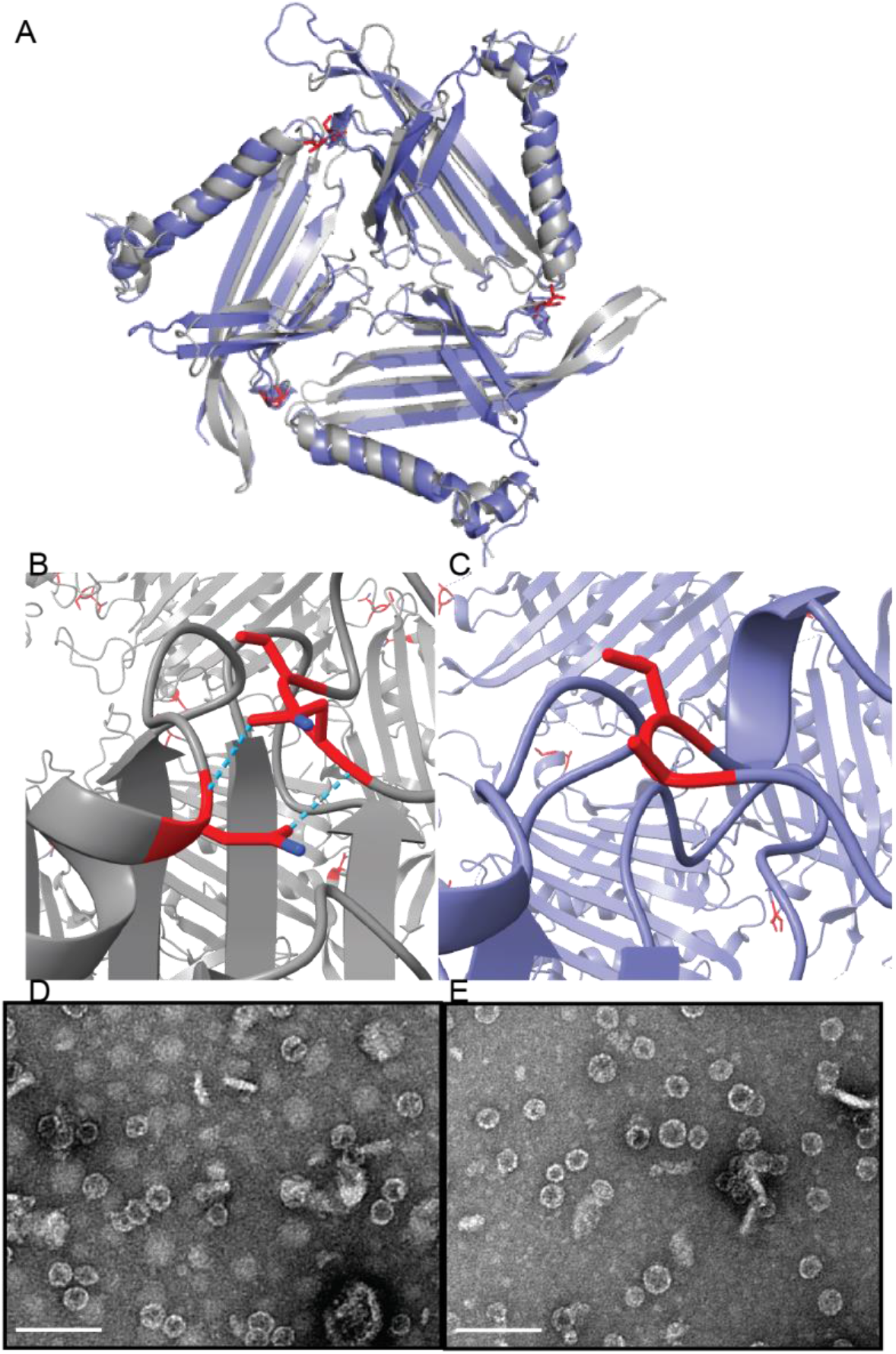
Qβ VLPs are structurally similar to MS2. A) Structural alignment between MS2 (PDB: 2MS2, gray) and Qβ (PDB: 1QBE, purple) B) Residues 36, 37, and 98 (red) in MS2 and hydrogen bonding interactions between them (blue) C) Residues 40 and 41 in Qβ (red). TEM images of D) Qβ WT VLPs with and E) Qβ A40P VLPs

### Generating and Assessing a 2D-AFL for Qβ 40/41

We wanted to assess if the trends we recorded in the MS2 36/37 2D-AFL were applicable in another VLP, so we again generated a 2D-AFL for residues 40 and 41 in Qβ (Figure 6A). Since the region in Qβ is not a precise match for the region in MS2, we reasoned that position 41 was an acceptable position to change when constructing a Qβ 2D-AFL. We completed similar quality checks as described earlier, and observed full coverage of the 441 mutants in the Qβ 2D-AFL. The average AFS of nonsense mutations was −3.2 for the Qβ 2D-AFL, further strengthening our assertion that the 40/41 Qβ 2D-AFL can be used to learn about Qβ VLP assembly.

**Figure 6:**
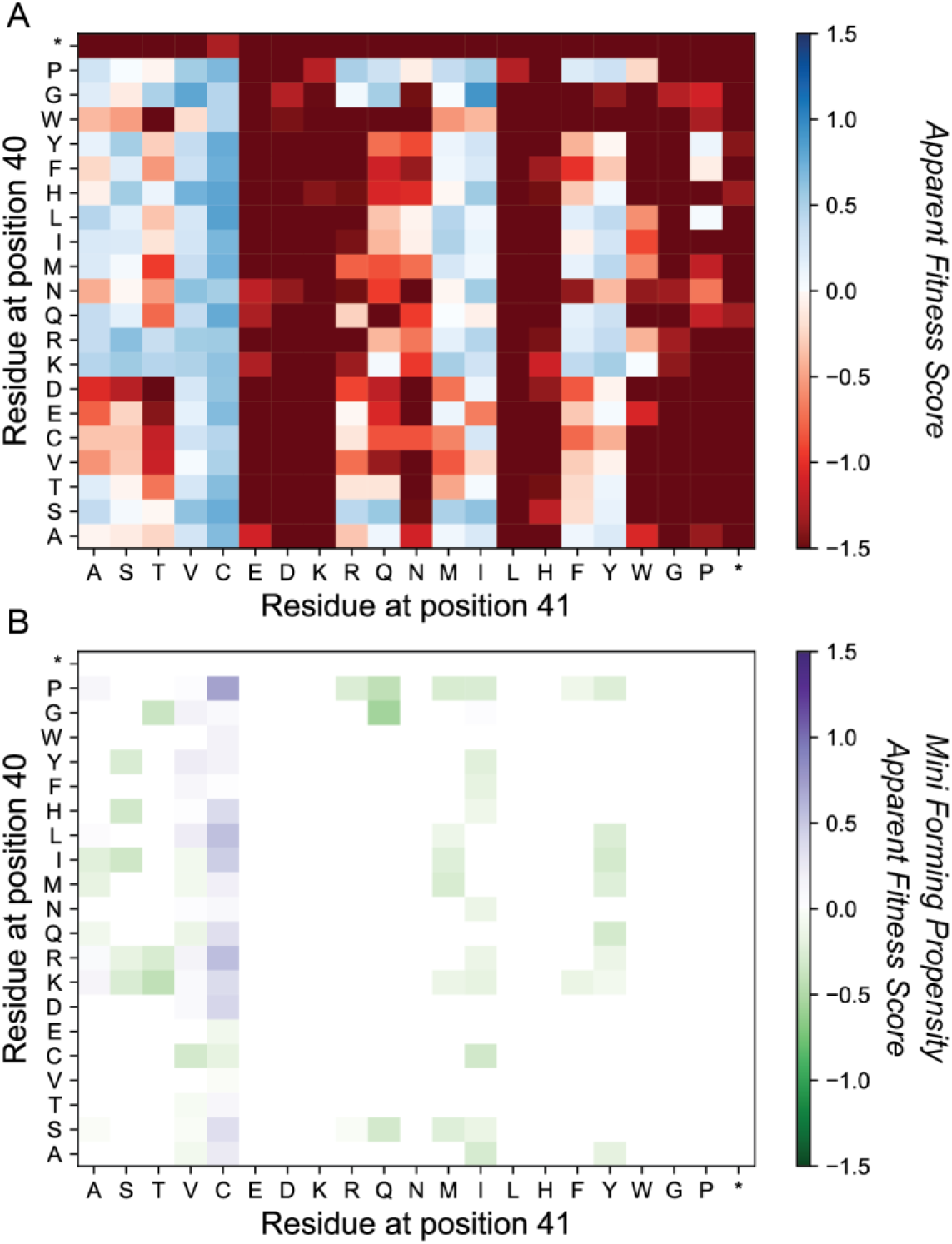
2D-AFL of Qβ 40/41 Library. A) 2D-AFL of the VLP selection library. B) Composite 2D-AFL where Mini Forming Propensity Apparent Fitness Scores are overlayed double variant combinations whose AFS >0.2.

We observed that in residue 40, mutations to serine, lysine, and proline have the highest number of AFS > 0.2 double mutants, but there was no banding trend like what we saw at residue 37 in MS2. However, there are bands of AFS > 0.2 double mutants for cysteine, isoleucine and valine mutations at position 41. Cysteine mutations appear to be an assembly tolerant single mutation in this region in both MS2 and Qβ VLPs. When we generated a composite Qβ 2D-AFL with MFP AFSs over positions in the 40/41 2D-AFL that have an AFS > 0.2, we saw that Qβ CP[V41C] containing double mutants have a trend of potentially forming smaller VLPs as shown in purple (Figure 6B). We performed similar HPLC SEC assembly validation assays for various 40/41 double mutants. As a negative control to see that our design heuristics applied to Qβ VLPs, we looked at three Qβ CP[V41I] mutants whose AFS > 0.2 but MFP AFS < 0.2 (predicted to form VLPs, but not Mini-sized): Qβ CP[A40T/V41I], Qβ CP[A40K/V41I], and Qβ CP[A40G/V41I]. HPLC SEC assembly validation of these individual mutants revealed that not only do the mutants with negative MFP AFSs only elute in WT characteristic elution range, but Qβ CP[A40G/V41I], MFP AFS = 0.01, also only elutes in the WT characteristic elution range (Supplemental Figure 5).

Since we established the 2D-AFL is functioning similarly to the MS2 36/37 2D-AFL, we wanted to validate if double mutants containing Qβ CP[V41C] formed “Mini” VLPs. The HPLC SEC assembly assay results indicated that both WT and Mini-sized VLPs formed (Figure 7A). The relative size bias scores (Figure 7B) for Qβ in this case seem predictive of only presence of smaller VLPs because the highest scoring double mutant Qβ CP[A40P/V41C] at 0.8 has both a WT and mini VLP characteristic peak.

**Figure 7:**
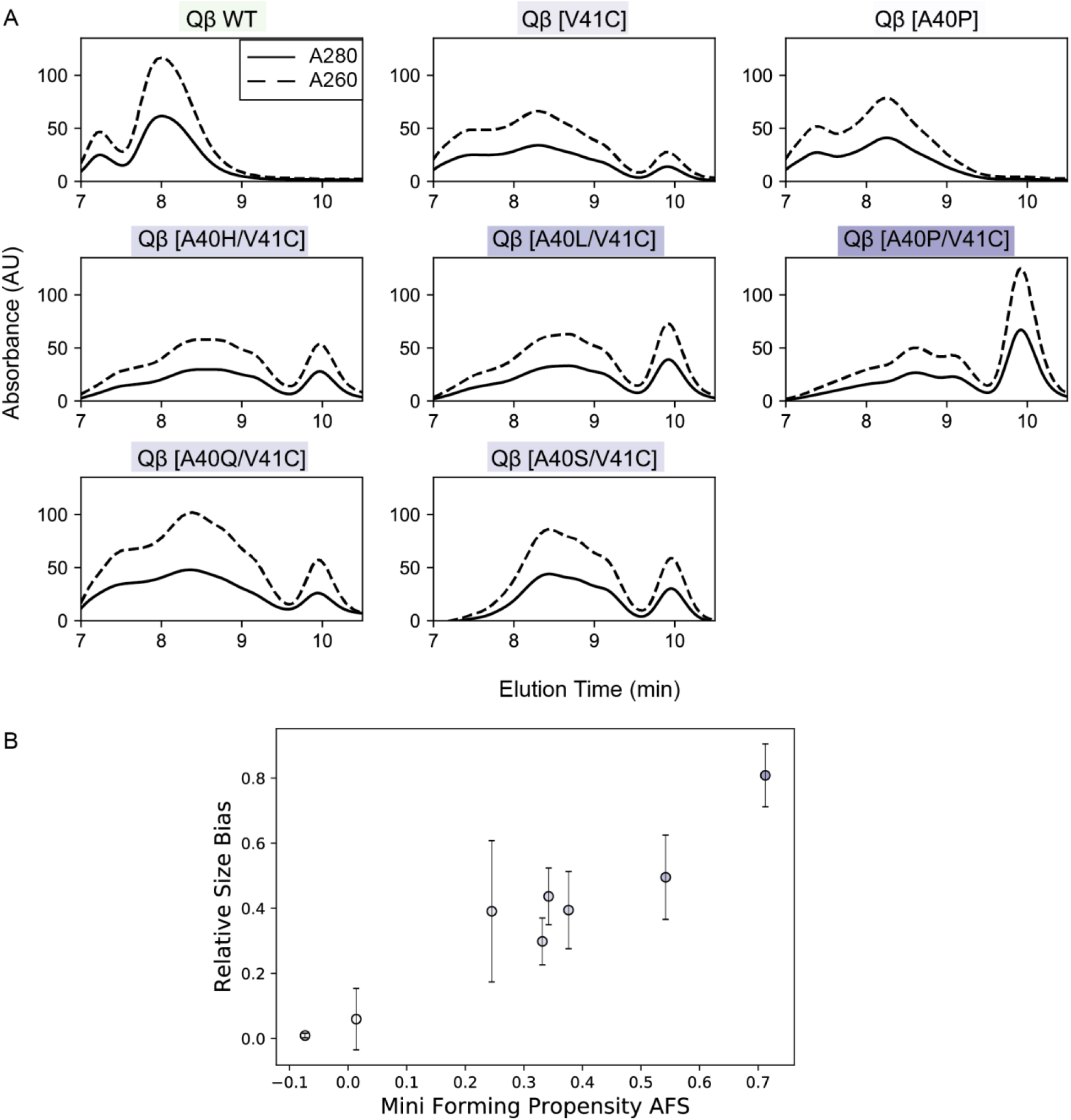
HPLC SEC Validation of Qβ 40/41 2D-AFL. A) HPLC Traces of double mutants that have AFS >0.2 and Mini Forming Propensity (MFP) AFS >0.2. Variant highlighted by MFP AFS value. B) Relative Size Bias plotted against Qβ MFP AFS

### Characterization of Qβ CP[V41C] Double Variant VLPs

Gaining more resolution on the structure of Qβ CP[V41C] containing variants would validate if the assembly metrics calculated predict biological Qβ VLP assembly phenomena. Qβ CP[V41C] was chosen first, to assess a baseline mutation that has not been previously assessed for a smaller sized phenotype. Two other double mutants, Qβ CP[A40L/V41C] and Qβ CP[A40P/V41C], were selected because they had AFS > 0.2, MFP AFS > 0.2, and the two highest relative size scores.

Like with the MS2 36/37 double mutant VLP characterizations, we visualized these variants with TEM and analyzed the distribution of sizes by counting at least 50 particles in each image (Figure 8). We see that the Qβ CP[V41C] CP variant forms a heterogeneous population of VLPs, similar to what we saw with Qβ CP[A40P] CP (Supplemental Figure 6). There is no proline to potentially shift away interactions with surrounding residues, so this result suggests that hydrogen bonding is not the only feature that could be involved with forming the smaller sized phenotype in Qβ VLPs. Qβ CP[A40L/V41C] and Qβ CP[A40P/V41C] also both form Mini and WT VLPs, but both variants also seem to form a prolate shaped VLP which forms as an elongated particle VLPs (Figure 8 C-D) [12].

**Figure 8:**
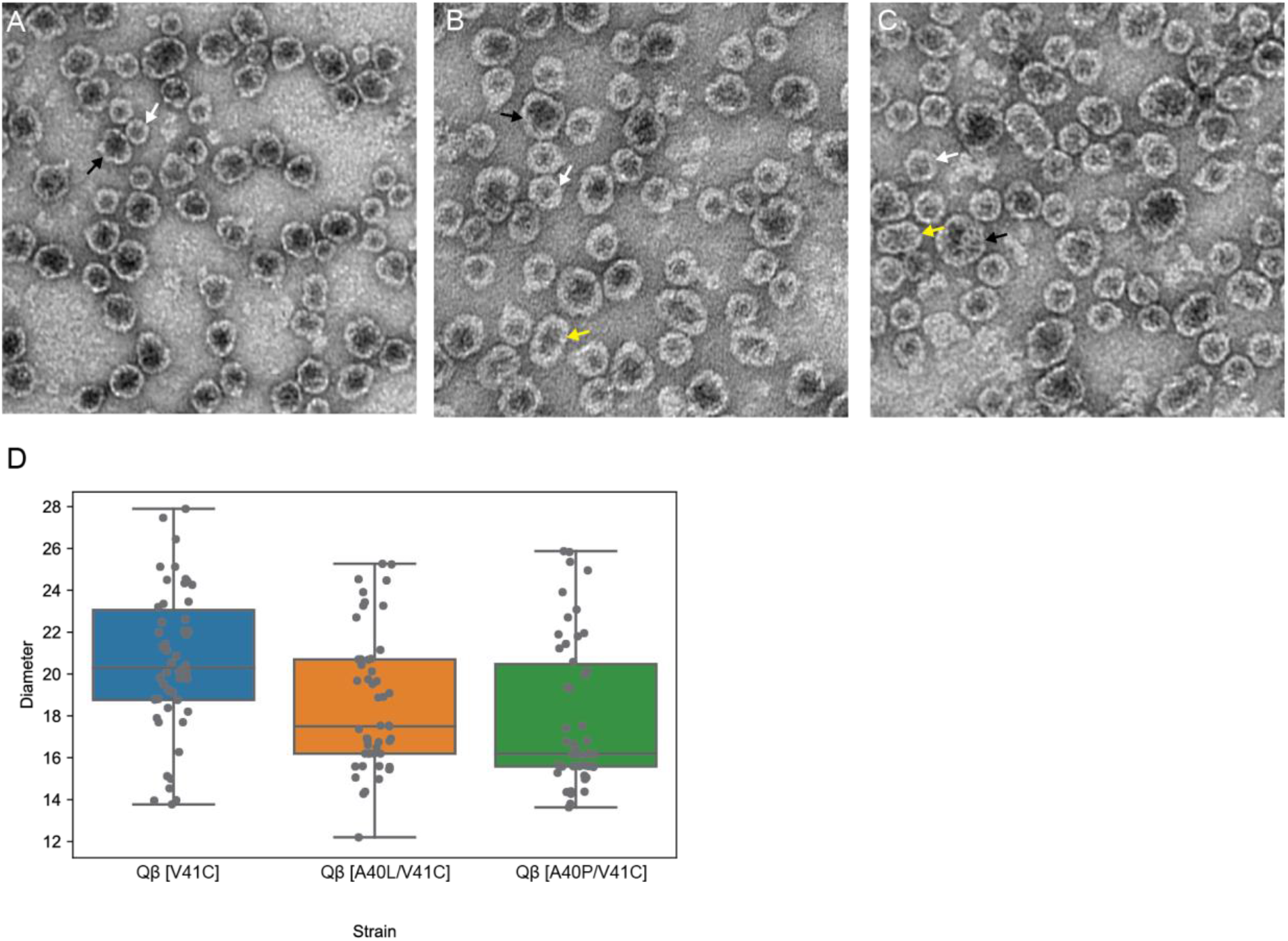
TEM Analysis of individual mutants for Qβ variants predicted to form Mini sized VLPs. White arrows point to mini VLPs, black arrows point to WT VLPs, and yellow arrows point to prolate VLPs. A) Qβ CP[V41C] B) Qβ CP[A40L/V41C] C) Qβ CP[A40P/V41C] D) Diameters of at least 50 VLPs quantified from TEM images are shown.

Qβ bacteriophages form small percentages of mini-sized assembled capsids naturally even if those variants are not infective and therefore are less than 5% of the particle population[12]. Crystal structure analysis of the VLP has shown that Qβ dimer conformations are flexible, being that the crucial VLP stabilizing interactions are intradimer salt bridges and other amino acid interactions. The robustness of their assembly was shown when “Mini” Qβ VLP variants were first identified with the Qβ CP [L35W] and Qβ CP [L35F][13]. These two Mini Qβ VLPs actually formed a heterogeneous population of VLPs, majorly forming small ~16.5 nm particles and less, but also forming 20 nm VLPs and longer prolate (23.5 × 20 nm) VLPs[13]. These alternate structure Mini Qβ VLPs showed that large amino acid variations at an interior facing residue disrupt WT VLP formation, but the flexibility of Qβ dimer conformations led to assembly of alternate geometry VLPs.

We have seen that variations to residues 40/41 of the Qβ CP in the same loop region as Mini Qβ variant and in analogous positions to the double mutant MS2 VLPs led to a heterogeneous size population dominated by smaller VLPs. We saw that Qβ CP[A40P], Qβ CP[V41C], Qβ CP [A40P/V41C], and Qβ CP[A40L/V41C] display consistent heterogeneous assembled VLP populations that were seen in Mini Qβ. The introduction of proline in this loop region in the Qβ CP conferred a similar smaller sized VLP phenotype as see in MS2, but the change in hydrogen bonding does not seem to be the defining property. Qβ CP[V41C] also confers the smaller sized particles, leading us to believe similar to Mini Qβ VLP variants, it was the disruption of potential intradimer side chain interactions that was a defining characteristic of forming smaller Qβ VLPs.

## Conclusions and Future Work

We have seen variations in analogous positions in two VLPs, that were not particularly sequenced aligned, led to a “mini” VLP formation. The characterization of our double mutants show that side chain interactions are crucial to smaller VLP formation, but the residue position themselves might only be predictive for VLPs that are stabilized by interdimer side chain interactions like in MS2 VLPs. Future work for these studies would be to crystallize double mutant-containing MS2 and Qβ VLPs and assess what sidechain interactions are present in the assembly and if these double variants lead to an increased number of conformations the CP can access.

## Acknowledgements

Portions of this project were funded by National Science Foundation Award CBET-2043973 to D.T.E.. E.C.H. was supported under by the Department of Defense, Air Force Office of Scientific Research, National Defense Science and Engineering Graduate (NDSEG) Fellowship (32 CFR 168a).

We would also like to acknowledge our collaborators in the Francis Lab and Lisa Burdette for helpful advice on this project.

