## Supplemental Information for "Systematic Engineering of Virus-Like Particles to Identify Self-Assembly Rules for Shifting Particle Size"

Supplemental Table 1: Plasmids used in this study.

| **Strain** | **Genotype** | **Antibiotic Resistance** |
| --- | --- | --- |
| BCI55 | pBAD MS2 WT | Chloramphenicol |
| BCI56 | pBAD MS2 [S37P] (Mini MS2) | Chloramphenicol |
| BCI54 | pBAD Non-assmebling MS2 WT | Chloramphenicol |
| BCI12 ^15^ | pBAD MS2 Golden-gate Entry Vector 2 | Chloramphenicol |
| BCI35 | pBAD MS2 [N36V/S37P] | Chloramphenicol |
| BCI39 | pBAD MS2 [N36I/S37P] | Chloramphenicol |
| BCI40 | pBAD MS2 [N36G/S37P] | Chloramphenicol |
| BCI73 | pBAD MS2 [N36G/S37C] | Chloramphenicol |
| BCI74 | pBAD MS2 [N36G/S37A] | Chloramphenicol |
| BCI77 | pBAD MS2 [N36E/S37F] | Chloramphenicol |
| BCI116 | pBAD MS2 [N36/S37P] | Chloramphenicol |
| BCI119 | pBAD MS2 [N36W/S37P] | Chloramphenicol |
| BCI120 | pBAD MS2 [N36M/S37P] | Chloramphenicol |
| BCI122 | pBAD MS2 [N36T/S37P] | Chloramphenicol |
| BCI123 | pBAD MS2 [N36P/S37P] | Chloramphenicol |
| BCI124 | pBAD MS2 [N36L/S37P] | Chloramphenicol |
| BCI125 | pBAD MS2 [N36Q/S37P] | Chloramphenicol |
| BCI28 | pBAD Qβ WT | Chloramphenicol |
| BCI148 | pBAD Qβ Golden-gate Entry Vector 2 | Chloramphenicol |
| BCI16 | pBAD Qβ [A40P] | Chloramphenicol |
| BCI87 | pBAD Qβ [A40H/V41C] | Chloramphenicol |
| BCI88 | pBAD Qβ [A40L/V41C] | Chloramphenicol |
| BCI89 | pBAD Qβ [A40T/V41I] | Chloramphenicol |
| BCI90 | pBAD Qβ [A40K/V41I] | Chloramphenicol |
| BCI91 | pBAD Qβ [A40Q/V41C] | Chloramphenicol |
| BCI94 | pBAD Qβ [A40P/V41C] | Chloramphenicol |
| BCI95 | pBAD Qβ [A40G/V41I] | Chloramphenicol |
| BCI98 | pBAD Qβ [A40L/V41S] | Chloramphenicol |
| BCI99 | pBAD Qβ [A40S/V41C] | Chloramphenicol |
| BCI128 | pBAD Qβ [V41C] | Chloramphenicol |

Supplemental Table 2: Primers used in this study

.

| **Primer** | **Description/Amplicon** | **Sequence (5’ to 3’)** |
| --- | --- | --- |
| ECH_DIS_5_3 | forward for A40P QuikChange Qβ | TCACAAGCGGGTCCAGTTCCTGCGCTGGAGAAGC |
| ECH_DIS_5_4 | reverse for A40P QuikChange Qβ | CAGCGCAGGAACTGGACCCGCTTGTGAAAGCGAG |
| ECH_DIS_5_9 | 36-37 NNK-NNK substitution using EMPIRIC cloning | aGGTCTCaCGCTaacggggtcgctgaatggatcagctctNNKNNKcgttcacaggcttacaaagtaacctgtagcgttcgtcagagctctGCGCtGAGACCt |
| ECH_DIS_5_10 | Forward primer with primer ECH_DIS_5_9 | aGGTCTCacgct |
| ECH_DIS_5_11 | Reverse primer with primer ECH_DIS_5_9 | aGGTCTCaGCGC |
| ECH_DIS_2_130 | F Amplicon Primer #1 (RNASeq PCR1) | TCGTCGGCAGCGTCAGATGTGTATAAGAGACAGgctcttaaagaggagaaaggtcatg |
| ECH_DIS_2_131 | R Amplicon Primer #1 (RNASeq PCR1) | GTCTCGTGGGCTCGGAGATGTGTATAAGAGACAGgccaaaacagccaagctttta |
| prBI192 | Index 1 N701 | CAAGCAGAAGACGGCATACGAGATtcgccttaGTCTCGTGGGCTCGG |
| prBI193 | Index 1 N702 | CAAGCAGAAGACGGCATACGAGATctagtacgGTCTCGTGGGCTCGG |
| prBI194 | Index 1 N703 | CAAGCAGAAGACGGCATACGAGATttctgcctGTCTCGTGGGCTCGG |
| prBI195 | Index 1 N704 | CAAGCAGAAGACGGCATACGAGATgctcaggaGTCTCGTGGGCTCGG |
| prBI196 | Index 1 N705 | CAAGCAGAAGACGGCATACGAGATaggagtccGTCTCGTGGGCTCGG |
| prBI197 | Index 1 N706 | CAAGCAGAAGACGGCATACGAGATcatgcctaGTCTCGTGGGCTCGG |
| prBI198 | Index 1 N707 | CAAGCAGAAGACGGCATACGAGATgtagagagGTCTCGTGGGCTCGG |
| prBI186 | Index 2 S501 | AATGATACGGCGACCACCGAGATCTACACtagatcgcTCGTCGGCAGCGTC |
| prBI187 | Index 2 S502 | AATGATACGGCGACCACCGAGATCTACACctctctatTCGTCGGCAGCGTC |
| prBI188 | Index 2 S503 | AATGATACGGCGACCACCGAGATCTACACtatcctctTCGTCGGCAGCGTC |
| prBI189 | Index 2 S504 | AATGATACGGCGACCACCGAGATCTACACagagtagaTCGTCGGCAGCGTC |
| prBI001 | Reverse for 36/37 NNK/NNK individual mutant golden gate inserts | aGGTCTCaGCGCagagctctgacgaacgctacaggttactttgtaagcctgtgaacg |
| prBI006 | MS2 [N36V/S37P] insert Forward | aGGTCTCaCGCTaacggggtcgctgaatggatcagctctGTTCCGcgttcacaggcttacaaagtaacc |
| prBI004 | MS2 [N36I/S37P] insert Forward | aGGTCTCaCGCTaacggggtcgctgaatggatcagctctATTCCGcgttcacaggcttacaaagtaacc |
| prBI021 | MS2 [N36G/S37P] insert Forward | aGGTCTCaCGCTaacggggtcgctgaatggatcagctctGGTCCGcgttcacaggcttacaaagtaacc |
| prBI223 | MS2 [N36G/S37C] insert Forward | aGGTCTCaCGCTaacggggtcgctgaatggatcagctctGGTTGTcgttcacaggcttacaaagtaacctgtagcgttcgtcagagctctGCGCtGAGACCt |
| prBI224 | MS2 [N36G/S37G] insert Forward | aGGTCTCaCGCTaacggggtcgctgaatggatcagctctGGTGGTcgttcacaggcttacaaagtaacctgtagcgttcgtcagagctctGCGCtGAGACCt |
| prBI225 | MS2 [N36G/S37A] insert Forward | aGGTCTCaCGCTaacggggtcgctgaatggatcagctctGGTGCTcgttcacaggcttacaaagtaacctgtagcgttcgtcagagctctGCGCtGAGACCt |
| prBI226 | MS2 [N36D/S37H] insert Forward | aGGTCTCaCGCTaacggggtcgctgaatggatcagctctGATCATcgttcacaggcttacaaagtaacctgtagcgttcgtcagagctctGCGCtGAGACCt |
| prBI006 | MS2 [N36E/S37F] insert Forward | aGGTCTCaCGCTaacggggtcgctgaatggatcagctctGAATTTcgttcacaggcttacaaagtaacctgtagcgttcgtcagagctctGCGCtGAGACCt |
| prBI018 | MS2 [N36C/S37P] insert Forward | aGGTCTCaCGCTaacggggtcgctgaatggatcagctctTGTCCGcgttcacaggcttacaaagtaacc |
| prBI019 | MS2 [N36W/S37P] insert Forward | aGGTCTCaCGCTaacggggtcgctgaatggatcagctctTGGCCGcgttcacaggcttacaaagtaacc |
| prBI008 | MS2 [N36P/S37P] insert Forward | aGGTCTCaCGCTaacggggtcgctgaatggatcagctctCCTCCGcgttcacaggcttacaaagtaacc |
| prBI009 | MS2 [N36T/S37P] insert Forward | aGGTCTCaCGCTaacggggtcgctgaatggatcagctctACTCCGcgttcacaggcttacaaagtaacc |
| prBI005 | MS2 [N36M/S37P] insert Forward | aGGTCTCaCGCTaacggggtcgctgaatggatcagctctATGCCGcgttcacaggcttacaaagtaacc |
| prBI003 | MS2 [N36L/S37P] insert Forward0 | aGGTCTCaCGCTaacggggtcgctgaatggatcagctctCTGCCGcgttcacaggcttacaaagtaacc |
| prBI003 | MS2 [N36Q/S37P] insert Forward | aGGTCTCaCGCTaacggggtcgctgaatggatcagctctCAACCGcgttcacaggcttacaaagtaacc |
| prBI242 | Gibson Primer Forward. Qβ EV2 backbone | ACTGTACAAATGAGGTCTCaCCTTCTCGCAATCGTAAGAA |
| prBI243 | Gibson Primer Reverse. Qβ EV2 backbone | TCACTGATAGGGAGGTCTCaATTTACCCCACGCGGATTGA |
| prBI273 | Gibson Primer Forward. Qβ EV2 GFP insert | aGGTCTCaAAATCCCACTAACGGCGTTGCCTCGCTTTCACAAGCG |
| prBI274 | Gibson Primer Reverse. Qβ EV2 GFP insert | aGGTCTCaAAGGCTGAGATACCGAAACGGTAACACGCTTCTCCAGCGCAGG |
| prBI244 | Reverse for 40/41 NNK/NNK Qβ inserts | aGGTCTCaGCGCCTGAGATACCGAAACGGTAACACGCTTCTCCAGCGCAGG |
| Qβ 40NNK/41NNK | 40/41 NNK/NNK Qβ geneblock | GGTCTCaCATGGCAAAATTAGAGACTGTTACTTTAGGTAACATCGGGAAAGATGGAAAACAAACTCTGGTCCTCAATCCGCGTGGGGTAAATCCCACTAACGGCGTTGCCTCGCTTTCACAAGCGGGTnnknnkCCTGCGCTGGAGAAGCGTGTTACCGTTTCGGTATCTCAGCCTTCTCGCAATCGTAAGAACTACAAGGTCCAGGTTAAGATCCAGAACCCGACCGCTTGCACTGCAAACGGTTCTTGTGACCCATCCGTTACTCGCCAGGCATATGCTGACGTGACCTTTTCGTTCACGCAGTATAGTACCGATGAGGAACGAGCTTTTGTTCGTACAGAGCTTGCTGCTCTGCTCGCTAGTCCTCTGCTGATCGATGCTATTGATCAGCTGAACCCAGCGTATTAAatGAGACCt |
| prBI256 | Qβ EV2 40G; 41I insert | aGGTCTCaCGCTCCCACTAACGGCGTTGCCTCGCTTTCACAAGCGGGTggtattCCTGCGCTGGAGAAGCGTGTTACCGTTTCGGTATCTCAGGCGCtGAGACCt |
| prBI257 | Qβ EV2 40L; 41S insert | aGGTCTCaCGCTCCCACTAACGGCGTTGCCTCGCTTTCACAAGCGGGTctgtctCCTGCGCTGGAGAAGCGTGTTACCGTTTCGGTATCTCAGGCGCtGAGACCt |
| prBI258 | Qβ EV2 40S; 41C insert | aGGTCTCaCGCTCCCACTAACGGCGTTGCCTCGCTTTCACAAGCGGGTtcttgtCCTGCGCTGGAGAAGCGTGTTACCGTTTCGGTATCTCAGGCGCtGAGACCt |
| prBI259 | Qβ EV2 40H; 41C insert | aGGTCTCaCGCTCCCACTAACGGCGTTGCCTCGCTTTCACAAGCGGGTcattgtCCTGCGCTGGAGAAGCGTGTTACCGTTTCGGTATCTCAGGCGCtGAGACCt |
| prBI260 | Qβ EV2 40L; 41C insert | aGGTCTCaCGCTCCCACTAACGGCGTTGCCTCGCTTTCACAAGCGGGTctgtgtCCTGCGCTGGAGAAGCGTGTTACCGTTTCGGTATCTCAGGCGCtGAGACCt |
| prBI261 | Qβ EV2 40T; 41I insert | aGGTCTCaCGCTCCCACTAACGGCGTTGCCTCGCTTTCACAAGCGGGTactattCCTGCGCTGGAGAAGCGTGTTACCGTTTCGGTATCTCAGGCGCtGAGACCt |
| prBI262 | Qβ EV2 40K; 41I insert | aGGTCTCaCGCTCCCACTAACGGCGTTGCCTCGCTTTCACAAGCGGGTaaaattCCTGCGCTGGAGAAGCGTGTTACCGTTTCGGTATCTCAGGCGCtGAGACCt |
| prBI263 | Qβ EV2 40Q; 41C insert | aGGTCTCaCGCTCCCACTAACGGCGTTGCCTCGCTTTCACAAGCGGGTcaatgtCCTGCGCTGGAGAAGCGTGTTACCGTTTCGGTATCTCAGGCGCtGAGACCt |
| prBI264 | Qβ EV2 40P; 41C insert | aGGTCTCaCGCTCCCACTAACGGCGTTGCCTCGCTTTCACAAGCGGGTccgtgtCCTGCGCTGGAGAAGCGTGTTACCGTTTCGGTATCTCAGGCGCtGAGACCt |
| prBI265 | Qβ EV2 40R; 41C insert | aGGTCTCaCGCTCCCACTAACGGCGTTGCCTCGCTTTCACAAGCGGGTcgttgtCCTGCGCTGGAGAAGCGTGTTACCGTTTCGGTATCTCAGGCGCtGAGACCt |
| prBI277 | Qβ EV2 40A; 41C insert | aGGTCTCaCGCTCCCACTAACGGCGTTGCCTCGCTTTCACAAGCGGGTgcttgtCCTGCGCTGGAGAAGCGTGTTACCGTTTCGGTATCTCAGGCGCtGAGACCt |

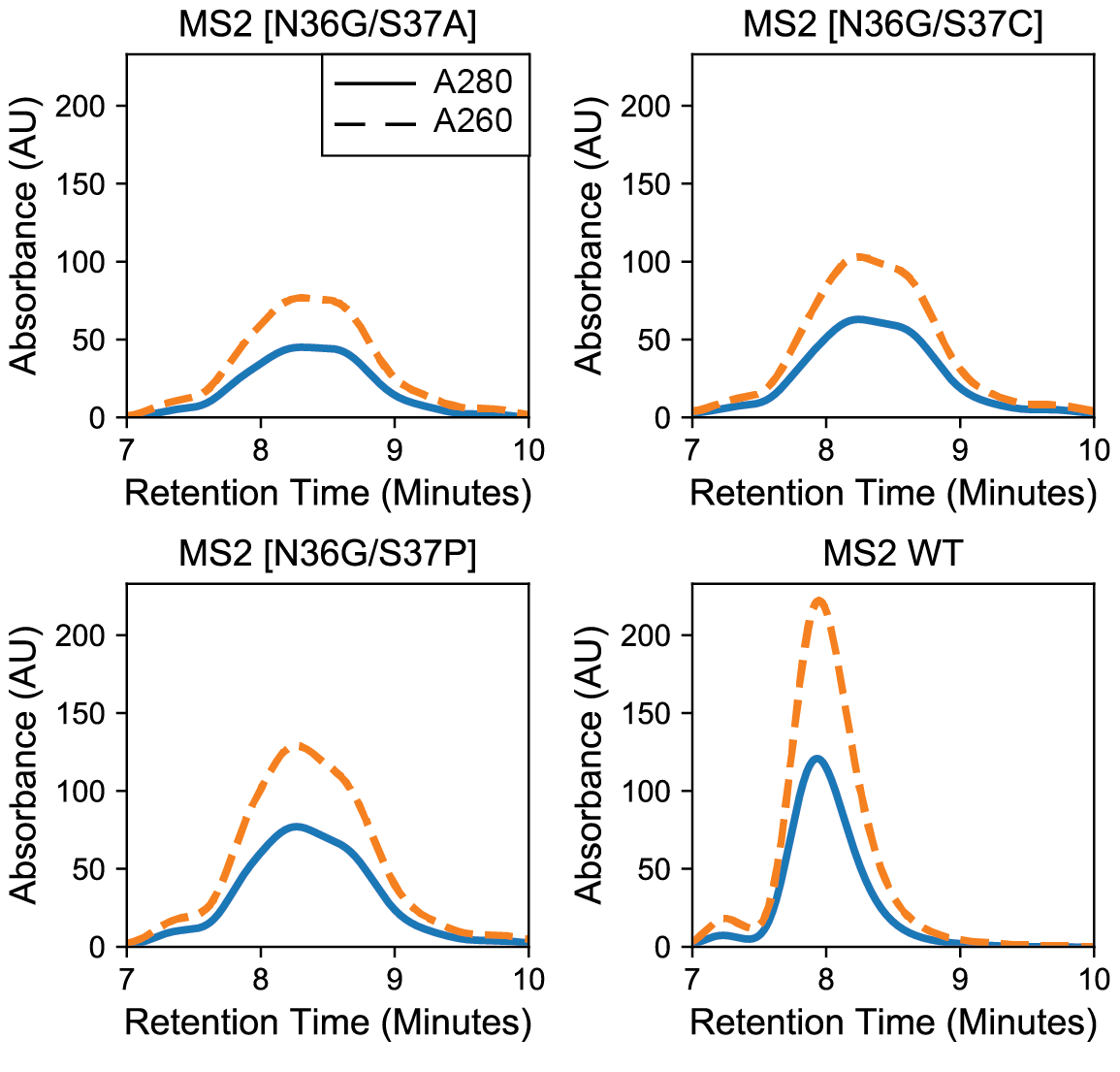

Supplemental Figure 1: HPLC SEC Traces for MS2 CP [N36G] containing double variants. *A) MS2 CP [N36G/S37A] B) MS2 CP [N36G/S37C] C) MS2 CP [N36C/S37A] D) MS2 CP WT*

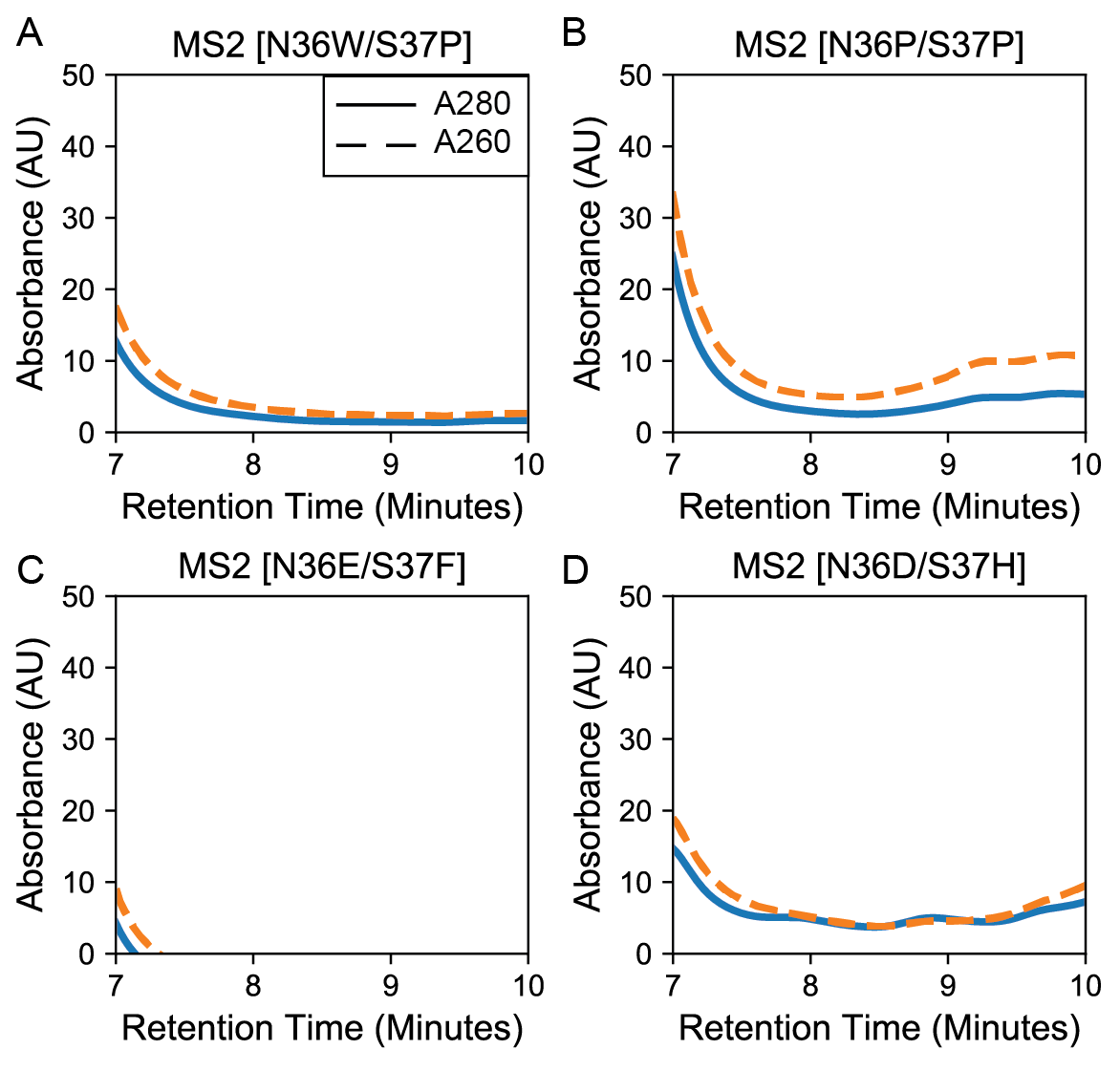

Supplemental Figure 2: HPLC SEC Traces for MS2 CP double variants whose AFS is negative or ~0 *A-B) MS2 CP double variants that have AFS <-0.2 C-D) MS2 CP double variants that have AFS ~0*

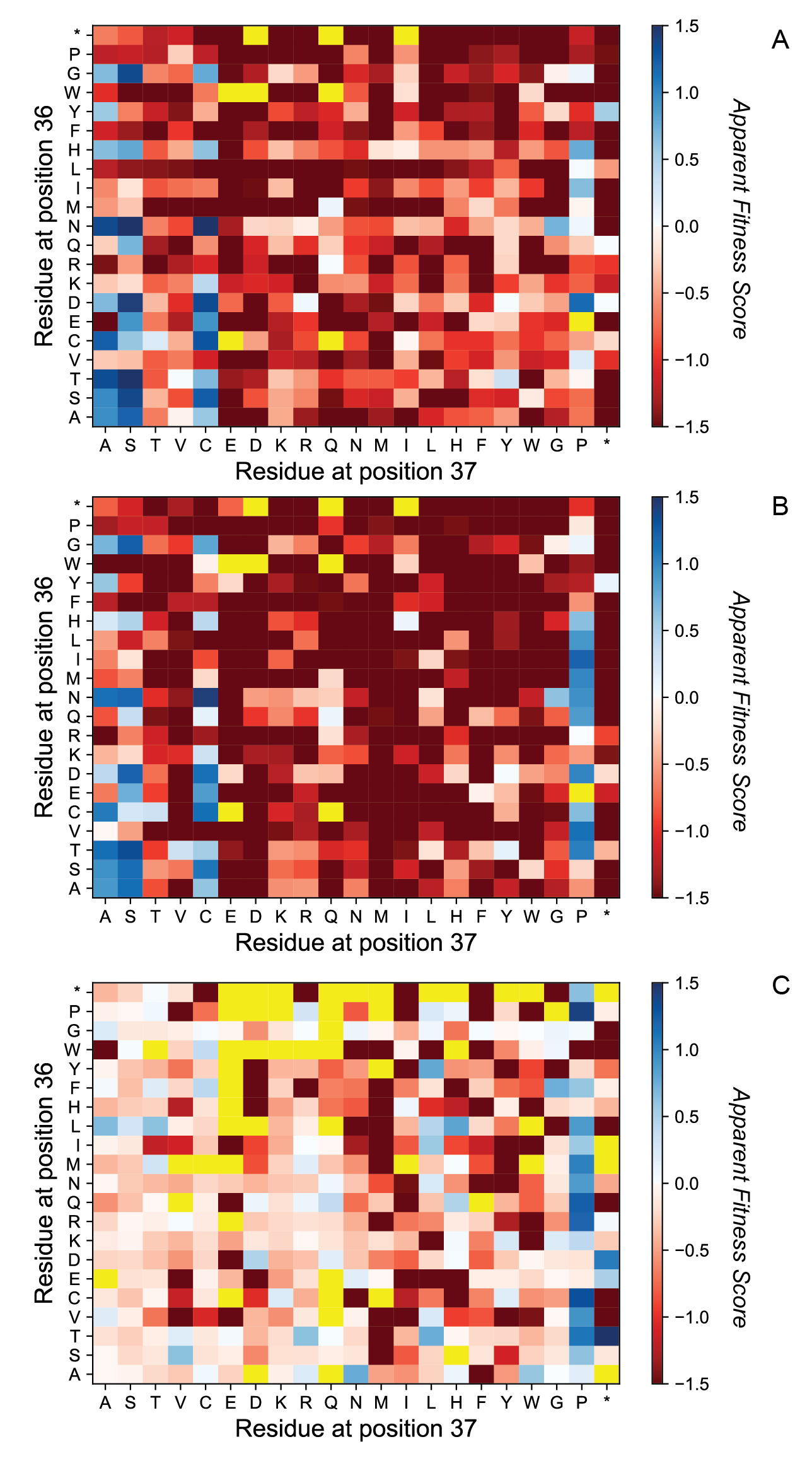

Supplemental Figure 3: 2D-Apparent Fitness Landscape of MS2 36/37 Library additional Assembly Fractions. *A) 2D-AFL of the Early Fraction VLP selection. B2D-AFL of the Late Fraction VLP selection. C) 2D-AFL of the Early Fraction/ Late Fraction. Blue indicates double variants were enriched after assembly selection and red indicates double variants that were less abundant after selection. Yellow indicates missing values.*

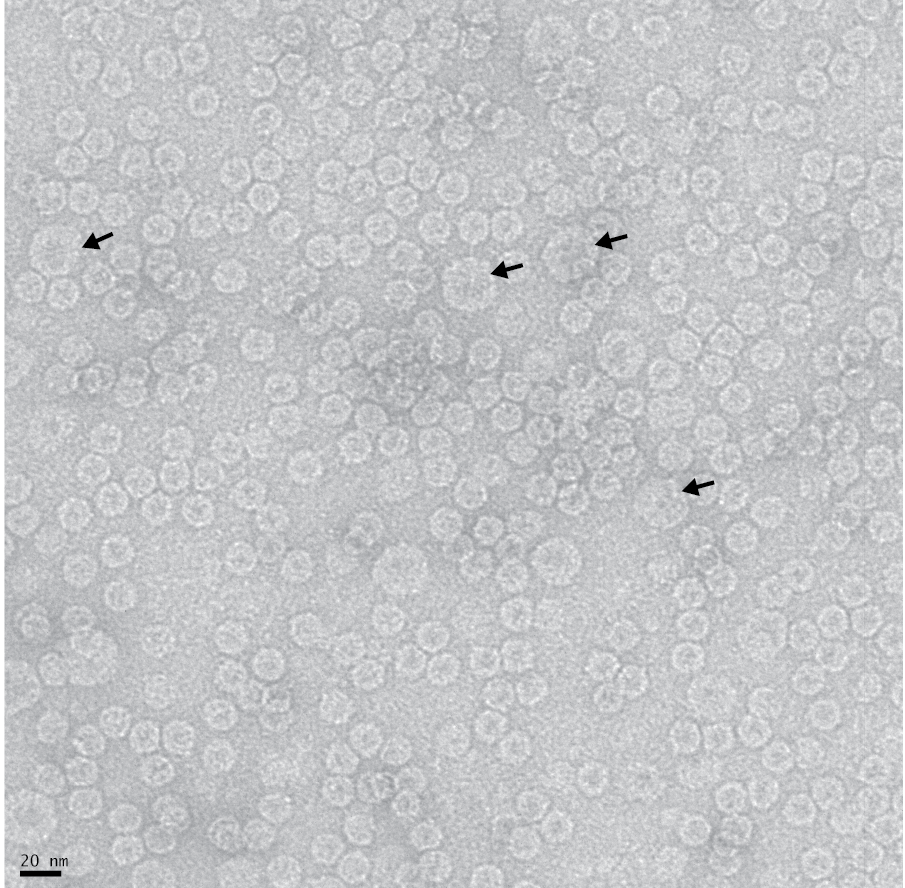

Supplemental Figure 4: TEM Analysis of WT characteristic elution HPLC fraction of MS2 CP[N36I/S37P]. *Black bar represents 20 nm. Black arrow points to WT sized VLPs*

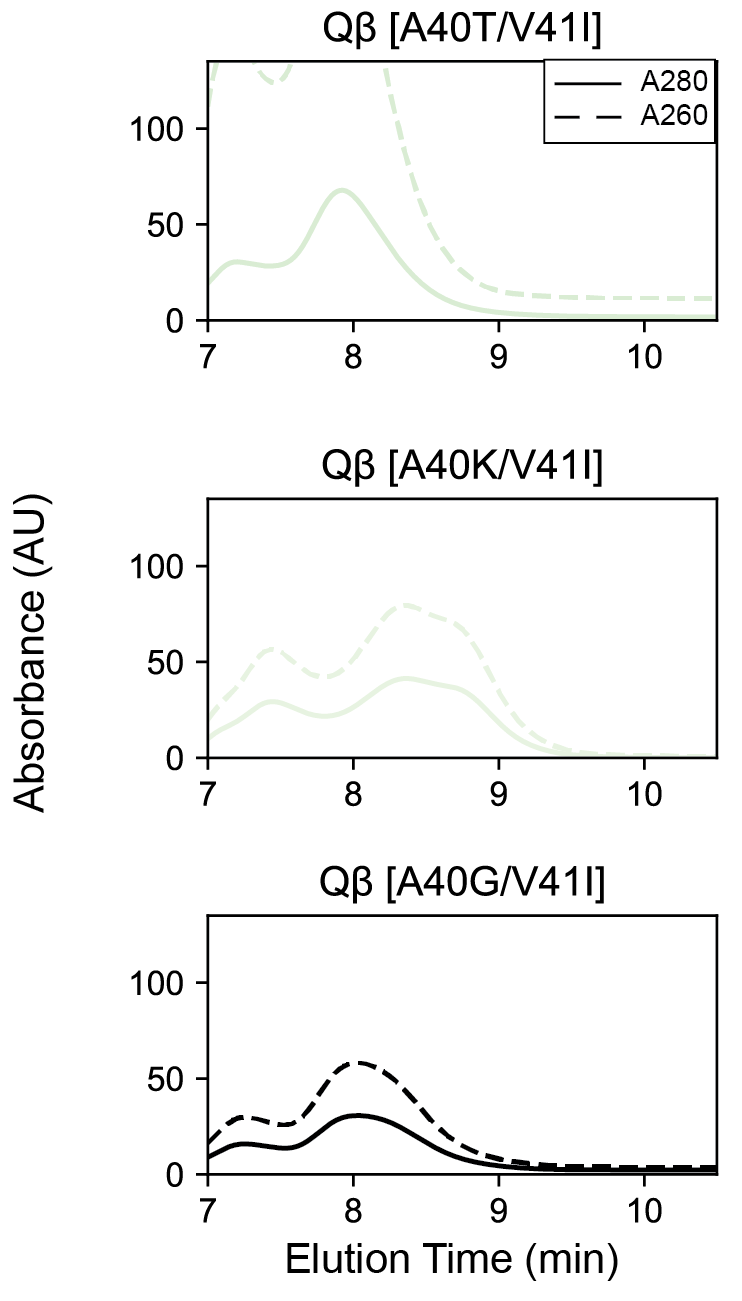

Supplemental Figure 5:HPLC SEC Traces for Qβ CP [V41I] containing double variants.

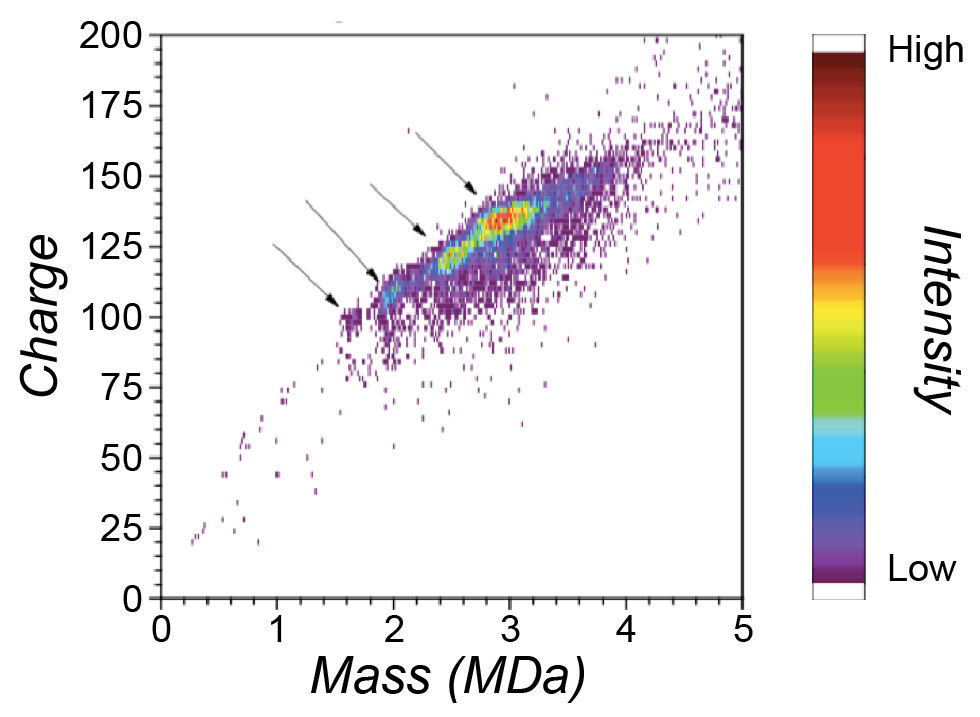

Supplemental Figure 6: Intact capsid mass spectrometry (CD-MS) indicates four populations of VLPs in purified Qβ A40P.
